# Collective radioresistance of T47D breast carcinoma cells is mediated by a Syncytin-1 homologous protein

**DOI:** 10.1101/448217

**Authors:** Roberto Chignola, Michela Sega, Barbara Molesini, Anna Baruzzi, Sabrina Stella, Edoardo Milotti

## Abstract

It is generally accepted that radiotherapy must target clonogenic cells, i.e., those cells in a tumour that have self-renewing potential. Focussing on isolated clonogenic cells, however, may lead to an underestimate or even to an outright neglect of the importance of biological mechanisms that regulate tumour cell sensitivity to radiation. We develop a new statistical and experimental approach to quantify the effects of radiation on cell populations as a whole. In our experiments, we change the proximity relationships of the cells by culturing them in wells with different shapes, and we find that the radiosensitivity of T47D human breast carcinoma cells in tight clusters is different from that of isolated cells. Molecular analyses show that T47D cells express a Syncytin-1 homologous protein (SyHP). We observe that SyHP translocates to the external surface of the plasma membrane of cells killed by radiation treatment. The data support the fundamental role of SyHP in the formation of intercellular cytoplasmic bridges and in the enhanced radioresistance of surviving cells. We conclude that complex and unexpected biological mechanisms of tumour radioresistance take place at the cell population level. These mechanisms may significantly bias our estimates of the radiosensitivity of breast carcinomas *in vivo* and thereby affect treatment plans, and they call for further investigations.

## Introduction

Breast cancer is the most common cancer in women worldwide, with 5-year survival rates that vary from 80% in developed countries to less than 40% in low-income countries [1]. Post-surgical adjuvant radiotherapy has been demonstrated to be effective in the control of local and regional microscopic residual disease and to reduce breast cancer-specific mortality, and also high-risk patients in the post-mastectomy settings benefit from radiotherapy [2, 3]. The positive outcome of radiotherapy for breast cancer is expected to improve further with the advancement of new radiotherapy techniques such as intensity-modulated radiotherapy, partial-breast irradiation and hypofractionation [3].

Quantitative predictions are required to calculate isoeffective radiation doses in alternative fractionation/protraction therapeutic schemes. Different mathematical models are used to this end. Their prediction capabilities, also within the settings of the novel radiotherapeutic approaches, are actively investigated and debated [4].

Model parameters are estimated by fitting model equations to experimental data and the problem is whether the experimental techniques return correct values or if they show limitations. This is very relevant in treatment planning, above all in the case of those tumours-such as breast tumours-that do benefit from radiation therapy.

The clonogenic assay is the common experimental approach to measure radiation sensitivity of tumour cells *in vitro* [4, 5]. After irradiation with different doses, cells are seeded in culture plates at appropriate dilutions to allow individual cell clones to proliferate and form colonies. Colonies grow, and within an incubation time of approximately two weeks they reach a size that is scored for growth. The number of positive colonies equals the number of cells surviving treatment. This simple experimental scheme has its drawbacks. First of all, not all cells in a tumour can originate a clonogenic progeny, a biological property shown only by cells with self-renewing potential (See e.g. refs.[6, 7] for an interesting discussion on this point). The fraction of such cells may be quite low [5], in the order of 10-30%, so that the effects of radiation are eventually measured only for a small fraction of cells in the tumour. Secondly, in a solid tumour clonogenic cells are not isolated and their proliferative potential is influenced by a tumour environment which includes non-clonogenic cells as well [8]. Indeed, tumours appear to be composed by hierarchically organized heterogeneous cell populations that orchestrate tumour progression [8], and it is known that the complex tissue organization of solid tumors also tunes the effects of radiation therapy [9, 10].

In our opinion, because of these drawbacks the standard clonogenic assay does not return a proper characterization of the radiation effects on tumour cell populations *in vivo.* We addressed this issue both statistically and experimentally. Here we present an approach that allows us draw robust statistical conclusions from radiation experiments with tumour cell populations. We use it to explore the biological basis of the collective radioresistance of breast carcinoma cells.

## Materials and methods

### Cell lines and reagents

Cells of the T47D human breast cancer cell line (ECACC, Salisbury, UK, number 85102201) were cultured at 37 °C in a humidified 5% CO2 atmosphere, in RPMI 1640 (Biochrom AG, Berlin, Germany) supplemented with 2 mM glutamine (Sigma-Aldrich, St.Louis, MO, USA), 35 mg/l gentamycin (Biochrom AG) and 10% heat-inactivated foetal bovine serum (Biochrom AG). We verified that the cell line used in this study was not cited in the Register of Misidentified Cell Lines maintained by the International Cell Line Authentication Committee (http://iclac.org/databases/cross-contaminations/). Cells were routinely tested and shown to be negative for mycoplasma contamination using MycoAlert™ PLUS Mycoplasma Detection Kit (Lonza Walkersville Inc., MD, USA,). Sytox AADvanced dead cell stain was provided by Invitrogen (Life Technology, Monza, Italy). Wheat Germ Agglutinin (WGA)-FITC and DAPI were from Sigma. Rhodamine phalloidin was from Cytoskeleton (Denver, CO, USA). We used the following antibodies (Ab): rabbit polyclonal Ab anti-HERV (ab 71115, Abcam); mouse monoclonal Ab anti-PActin (Sigma); goat anti-rabbit IgG-FITC (ab 6717, Abcam); goat anti-mouse IgG-FITC (ab 6785, Abcam).

### Irradiation

A Gammacell40 irradiator (Atomic Energy of Canada Limited, Kanata, Ontario, Canada) was used to treat cell cultures. The instrument is equipped with a ^137^Cs source and both the dose-rate and the dose-uniformity within the sample tray is routinely monitored by the Radiation Protection Service of the University of Verona. The dose rate was 0.6654 Gy/min. The measured uniformity was ± 1.3 % over the entire sample chamber.

### Clonogenic assay

Clonogenic assays were performed as described in [5] with minor modifications. Briefly, T47D cells from sub-confluent monolayer cultures were detached by trypsin treatment, counted using a hemocytometer and seeded in 6-well plates using standard medium enriched with 33% T47D conditioned growth medium. After 24 hours the cells were exposed to different doses of gamma-rays. For each cell dilution 4 replicates were performed. After 16 days the colonies were fixed with methanol and stained with Giemsa stain. Images were acquired using a desktop scanner (Epson Perfection V700 Photo; resolution settings: 1200 DPI) and analyzed using the ImageJ 1.47v software (NIH, USA; website: http://imagej.nih.gov/ij/). Colonies containing more than 50 cells were counted and scored positive for growth. Overall we counted 973 colonies with an average of 195 colonies for each treatment condition. Plating efficiency was determined dividing the number of counted colonies by the number of plated cells (plating efficiency of control samples was ~25%). The surviving fraction was calculated as the ratio of plating efficiency for irradiated and control samples. Data were fitted with the linear-quadratic model [4]:

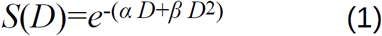

where *S(D)* is the surviving fraction of cells at a given dose of radiation *D* (Gy) and *a* and are non-negative parameters.

### Effects of radiation on cells: statistical model

Standard clonogenic assays measure how many clonogens, i.e., cells with self-renewing potential, survive a given dose of radiation. Let *S(D)* denote the survival probability of clonogens irradiated with dose *D* (eq. 1) and *ε* the fraction of clonoges in a population of *μ* cells on average. Then, the mean number of clonogens that survive irradiation is *S(D)sp* If cells do not interact with each other and/or with the other cells in the population, then the probability that they survive a given dose of radiation follows the Poisson distribution. Therefore, the probability that no clonogen survives is:

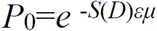

and the probability that at least one clonogen survives is:

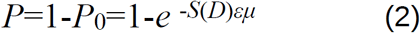

### Effects of radiation on cells: experiments

We seeded the cells into the wells, either flat-bottom or conical-bottom, of 96-well culture microplates at the indicated cell concentrations (see e.g. Fig.1) to grow independent cell populations (final volume 200 μl). After 24 hours, the plates were treated with gamma-rays at the dose of 8 Gy. Control sham-treated cell populations were kept in the irradiator room for the whole treatment duration. Microplates were then transferred into the CO2 incubator. After ~7 days (control populations) or ~40 days (irradiated populations), cells populations were scored as surviving or sterilized by careful microscopic analysis (carried out by three independent and blinded observers) and ATP measurements. ATP was measured by the luciferine/luciferase assay using the CellTiter-Glo luminescent cell viability kit (Promega, Milan, Italy). Luminescence was measured using a microplate luminometer (FLX800, Bio-Tek Instruments, Bad Friedrichshall, Germany) and the data expressed in luminescence arbitrary units. Background ATP levels were determined 48 hours after cell treatment with a lethal dose of 20 Gy. In long-lasting experiments we carefully checked for medium evaporation and we replenish the wells with fresh medium to keep the growth volume constant.

**Fig.1.**
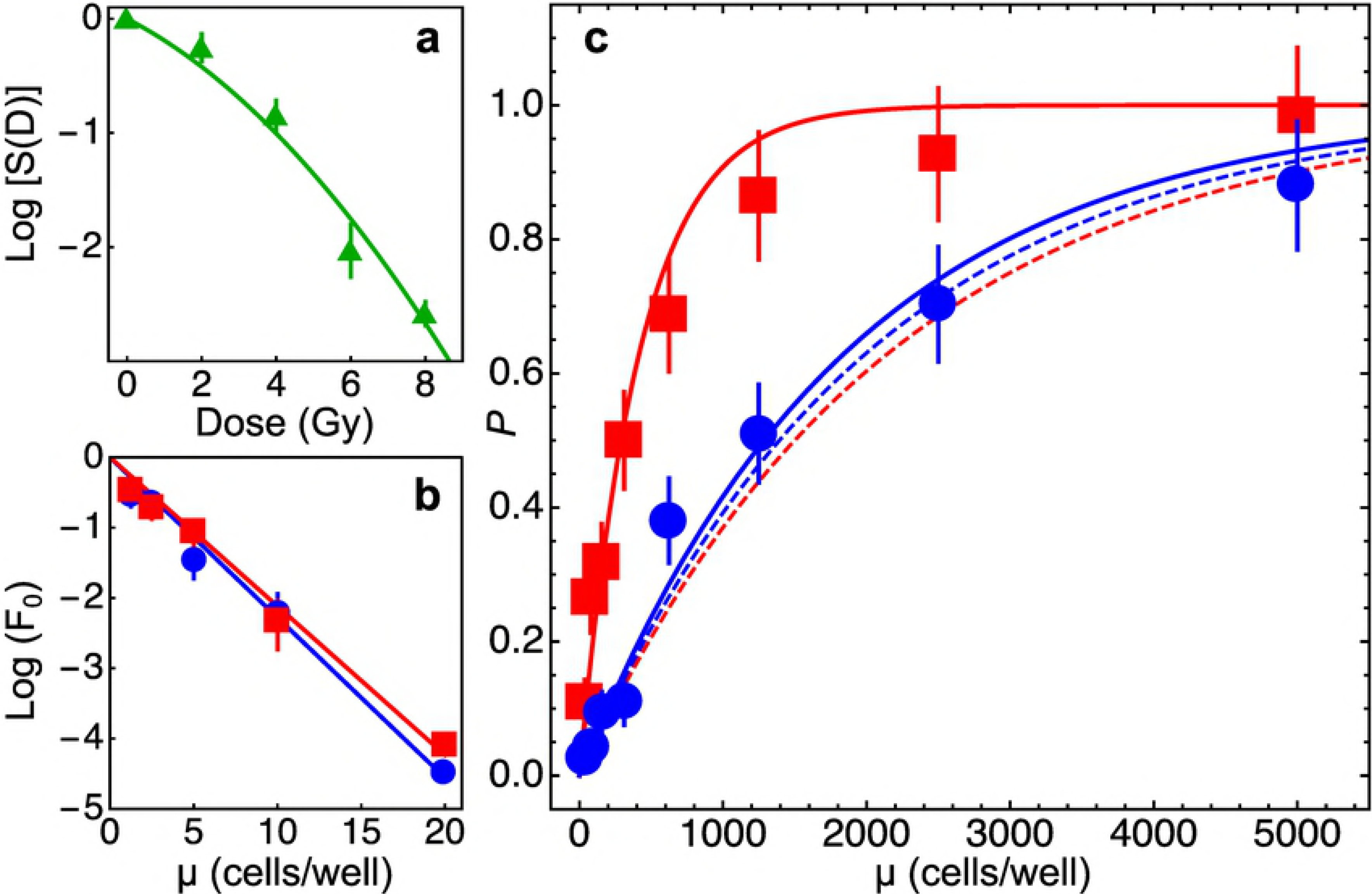
Radiosensitivity of T47D cells. Under the assumption that only clonogenic cells are radiosensitive, the survival probability of cell populations is given by (see Materials and Methods, eq.2) *P=1-Po=1-e*-^S(D)si^, where *S(D)* is the fraction of clonogenic cells surviving a dose *D* of radiation, *s* is the fraction of clonogenic cells in the population and *i* is the mean number of cells in the sample. **a:** estimate of *S(D)* using standard clonogenic assays. Data were fitted with the linear-quadratic model (eq.1, see Materials and methods): α=0.389±0.13 Gy^1^, y0=0.047±0.01 Gy^-2^, *α/β=8.22* Gy(*x*^2^/df=4.97). Thus S(8 Gy)~0.0022. **b:** estimate of *s* in limiting dilution assays with non irradiated cells. Cells were seeded either in F-bottom (blue circles and line) or V-bottom (red squares and line) wells at the indicated mean number *i* of cells/well and the fraction of wells showing no proliferation F_0_ was determined. Data were fitted with eq.3 (see Materials and methods): s=0.228±0.0097 (r^2^=0.99, cells in F-bottom wells), s=0.212±0.0081 (r^2^=0.99, cells in V-bottom wells). **c:** survival probability *P* of independent cells populations grown in F-bottom (blue circles and lines) or V-bottom (red squares and lines) wells starting from a mean initial number of cells/well *μ* and irradiated with a dose of 8 Gy. Dashed lines: model predictions with eq.2 and experimentally determined parameters. In this case the model can be exploited as the null hypothesis for statistical testing. Data from cell populations grown in F-bottom wells did not significantly deviate from model predictions (x^2^=6.69, p=0.35, goodness-of-fit test), whereas data obtained with cell populations grown in V-bottom wells did not agree with model predictions and the null hypothesis could be rejected (x^2^=489.4, p~10^-102^, goodness-of-fit test). Continuous lines: nonlinear fits of experimental data with eq.2. and with only S(D) as a free parameter. The estimated surviving fraction was ~4.7 fold higher for cell populations grown in V-bottom wells (S(D)=0.011±0.0012) than for cells grown in F-bottom wells (S(D)=0.00236±0.0002). Overall the data show that the sensitivity of cell populations to radiation treatment does not only depend on the number of clonogenic cells in the population, and that tumor cell populations grown under experimental conditions that favour cell contacts are more radioresistant.

### Limiting dilution assays

We measured the fraction of clonogenic cells in populations of cells seeded in flat-bottom and conical-bottom 96-wells microplates by limiting dilution assays [11]. Homogeneous single-cell suspensions were obtained by careful trypsin treatment of cell cultures. The cells were then randomly and independently distributed into the wells of 96-wells microplates, one plate for each cell concentration ranging from 1 to 20 cells/well. After ~30-40 days the wells showing absence of proliferating cell populations were counted. Under these conditions, the fraction of negative wells (the ratio between the number of negative wells and the total number of seeded wells) obeys Poisson statistics [11]:

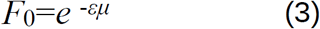

where *F*_o_ is the fraction of negative wells, *μ* the mean cell concentration expressed as number of initially seeded cells per well and *ε* is the fraction of clonogenic cells.

### Spheroid cultures and the outgrowth method

T47D spheroids were obtained by the liquid overlay method as described previously [12, 13]. Briefly, cells (500 cells/well in 150 μl of standard culture medium) were plated in 96 round-bottomed well plates previously coated with an agar gel layer (50 μl of a 0.7% w/v solution of agar in complete medium) to initially prevent cell attachment. Spheroids of about 400 pm diameter were transferred into the wells of 6-well culture plates (one spheroid/well in 2 ml culture medium). The wells were coated with a 2 ml layer of agar to grow spheroids in suspension. Spheroids of about similar size were isolated and transferred into the wells of 24-well culture plates (one spheroid/well in 1 ml culture medium) or in glass bottom Ibidi chambers (for confocal analyses, see below) and allowed to adhere to obtain rings of densely-packed cells outgrowing from spheroids. After 24 hours the cultures were exposed to a dose of 8 Gy of ionizing radiations.

### Microscopy

We used an Evos (AMG, Life Technology) digital inverted microscope to routinely monitor cell morphology. Cells were also analysed by confocal microscopy. To this purpose, we grew cell monolayers in glass bottom μ-Slide IbiTreat chambers (Ibidi GmBH, Martinsried, Germany) using the outgrowth method (see above). Cells were fixed with 4% (w/v) paraformaldehyde for 30 min and, after washings with PBS and quenching with 50 mM NH4G, permeabilized with PBS-0.1% Triton X-100. Cells were then stained with WGA-FITC (to label cytoplasmic and membrane glycoproteins containing β(1,4)-N-acetyl-D-glucosamine) and rhodamine-phalloidin (to label F-actin) for 30 min and then with DAPI (to label nuclei) for 10 min. Images at Az=1.76 pm were collected using the SP5 confocal microscope from Leica Microsystems (Mannheim, Germany) with 63x objective (HCX PL APO A blue 1.4NA OIL). Image analyses were performed with ImageJ 1.47v sofware.

### Flow cytometry

Cell surface and intracellular expression of biomarkers was investigated by flow cytometry. For cell surface analyses, the cells were kept on ice and incubated with primary antibodies for 30 min. After washings with cold PBS, the cells were incubated for further 30 min with secondary FITC-conjugated antibodies. Dead cells were labelled with the vital fluorescent dye Sytox. For intracellular analyses, the cells were permeabilised using the Fix-Perm kit from Nordic-MUbio (Susteren, The Netherlands) following the manufacturer’s instructions. We measured cell-associated fluorescence using a Guava easyCyte™ 5 flow cytometer, controlled by GuavaSoft v.2.7 software (Merck Millipore, Billerica, MA, U.S.A.). The cytometer was equipped with 488 nm, 20 mW, blue laser light, and forward scatter (FSC) photodiode and side scatter (SSC) photomultiplier. Green fluorescence 525/30 filter, yellow 583/26 and red 680/30 filters allow analysis of fluorescence emissions from samples. Calibration of the cytometer was routinely checked using the Guava EasyCheck kit (Merck Millipore, Billerica, MA, U.S.A.) according to the manufacturer’s instructions. Raw listmode data were exported as CSV files and then imported in the software Mathematica for data processing and analyses.

### Reverse Trascription PCR (RT-PCR) and sequence analysis

Total RNA was extracted from T47D mammary carcinoma cells using TRIzol reagent (Thermo Fisher Scientific). After DNase treatment, total RNA was further purified using ‘RNeasy Mini Kit’ (QIAGEN) following the RNA cleanup protocol. RNA quality was evaluated using Agilent 2100 Bioanalyzer (Agilent). Two micrograms of RNA were reverse-transcribed using ‘GoScript reverse transcription system kit’ (Promega) in a total volume of 20 pl according to the manufacturer’s instructions. The RT primer mix contained both oligo(dT)15 and random primers. Five microlitres of the complementary DNA (cDNA) were used as template for the subsequent PCR analysis with the following primers (F, 5′-AGCCCCGCAACAAAAGAGTA-3′ and R, 5′-TCCCGATCATTGTCCCTCC-3′). The amplified DNA fragment encompasses a region of 683 nt of the 3′ terminal coding region and a portion 276 nt of the 3′ untranslated region of the Syncytin-1 (AF208161) transcript. The resulting PCR product was further cloned into the pGEM®-T Easy Vector (Promega) and checked by sequencing. A homology search of the cDNA sequences was carried out using the National Center for Biotechnology Information website (http://www.ncbi.nlm.nih.gov/BlastN). Both the cDNA sequence and the deduced amino acid sequence were aligned to those of human Syncytin-1 using Clustal Omega software [14] hosted at https://www.ebi.ac.uk.

### Statistics

The reduced *x^2^* = *X^2^/df* where *df* is the number of degrees of freedom) was used to determine the goodness of the nonlinear fits. All mathematical and statistical analyses were carried out using the software Mathematica (Wolfram Research Inc., Champaign, IL, USA).

### Data availability

The authors declare that all data supporting the findings of this study are available from the corresponding author on reasonable request.

## Results

### Radiobiology of cell populations

The survival probability of cell populations composed of an average number of cells *i* to a given dose of radiation, under the assumption of statistical independence between cells, is given by eq.2 (see the Materials and methods section for details).

We estimated the values of both parameters *ε* and *S(D)* in independent assays (Fig.1), and eq.2 was used as the null hypothesis for goodness-of-fit statistical tests.

We seeded the cells into the wells of culture microplates with different well geometry, either flat-bottom (F) or conical-bottom (V), to vary the chance of the cells to keep contact with neighbouring cells. To estimate the survival probability *P* (eq.2), for both experimental conditions we grew 96 independent cell populations starting from a sample of *μ* cells, with *μ* ranging between 19.5 and 5000 cells/well (8 different cell density values), for an overall number of 1536 cell populations (Fig.1). Cell populations growing in V-bottom wells showed a higher probability to survive 8 Gy treatment with respect to those growing in F-bottom wells. The dose of 8 Gy was chosen on the basis of mathematical simulations carried out with eq.2 and experimentally determined parameters to find suitable experimental conditions linking together radiation dose, average cell numbers and survival probabilities.

The null hypothesis could be rejected for cell populations growing in V-bottom wells but not for those growing in F-bottom wells (Fig.1). Thus, a higher chance to establish contacts between cells in a population is correlated to a higher radioresistance of the population as a whole.

### Morphological analyses of irradiated cells

To start investigating the biological basis of the observed radioresistance of T47D cells when cultured in conical-bottom wells we first carried out morphological analyses with a particular type of monolayer cell cultures in which cells reach high densities per unit area. In traditional cell cultures, cells reach high density at confluence when, however, the environment becomes toxic because of nutrient deprivation, accumulation of waste molecules and reduced pH. To obtain confluent monolayer cultures suitable for long-lasting experiments we resorted to an experimental technique named the “spheroid outgrowth method” [15, 16]. First we grew three-dimensional cell cultures (spheroids) up to a diameter of ~400 μm, a suitable size to isolate the spheroids by micro-manipulation. Individual spheroids were then placed into the wells of 24-well culture plates and allowed to adhere to their plastic surface. Under these conditions, the cells progressively leave the spheroid cluster and form a ring of densely-packed proliferating cells around it. The ring contains up to a few hundred cells that grow in an extended nutrient-rich environment that can sustain cell growth for a long time. At the same time the ring of cells outgrowing spheroids can be easily analysed under the microscope.

After 96 hours from irradiation with 8 Gy, giant cells with multiple nuclei appeared in the outgrowing rings (Fig.2). Most cells became giant and multinucleated 144 hours after irradiation. Cells with diameters 3 times higher than those measured in control, sham-treated, samples were observed, and it was difficult to discriminate the individual cell bodies for cells located close to the adherent spheroid core. The heterogeneous morphology of the cell population was evident 336 hours after irradiation, and cytoplasmic bridges between cells could also be observed. Meanwhile the cells in the rings of irradiated spheroids did not proliferate (Fig.2). At the beginning of the post-irradiation regrowth phase (~500 hours, Fig.2) a few giant cells with multiple nuclei were still present in the population. However, the population was mostly composed by individual mononucleated cells likely deriving from the division of the giant cells. 600 hours after irradiation the cell population around the spheroid core proliferated again and the cells’ morphology was similar to that of control sham-treated cells (Fig.2).

**Fig.2.**
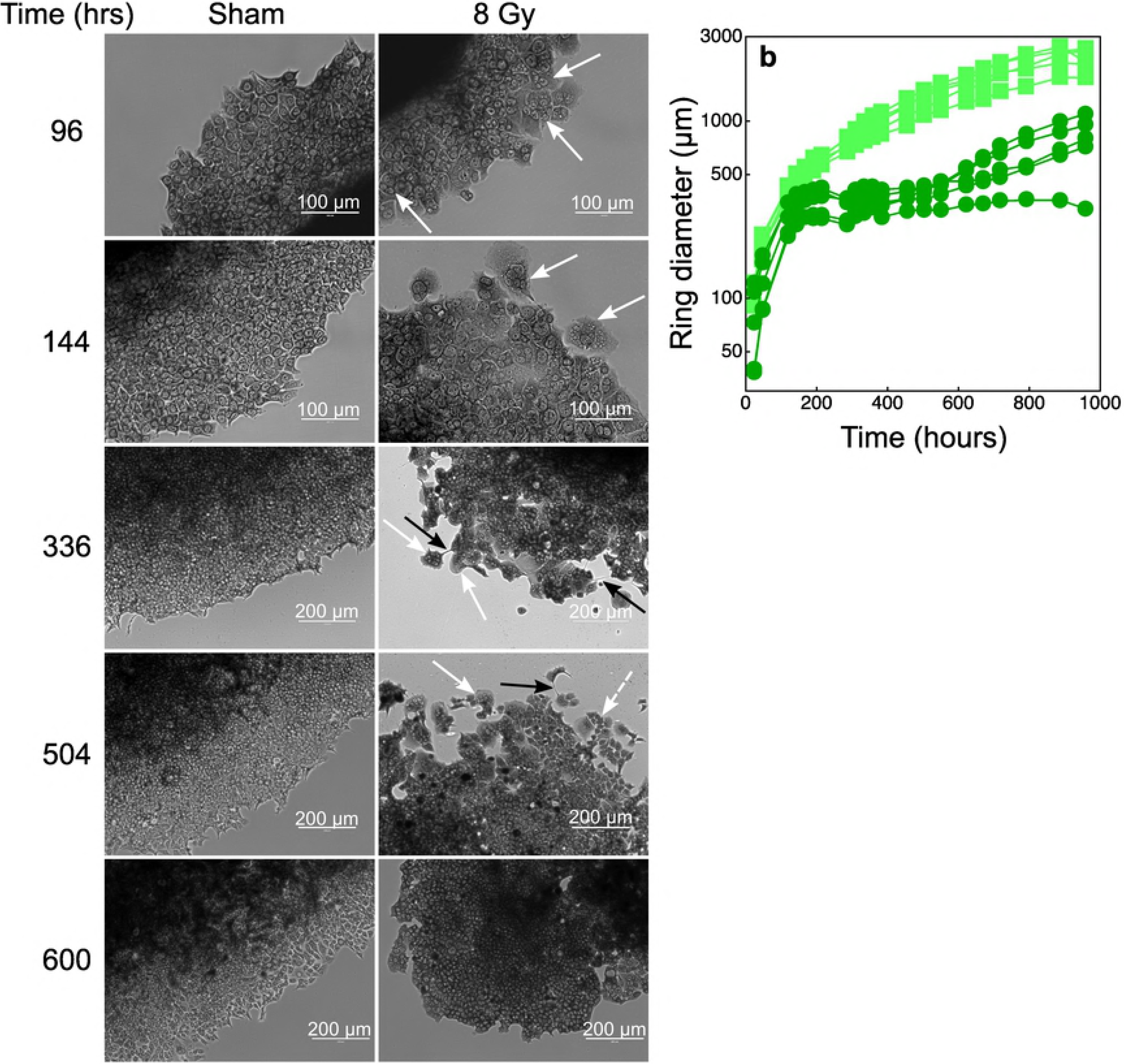
Cytological analyses of the cells in the rings around irradiated (8 Gy) and sham-irradiated tumour spheroids. The spheroid outgrowth method (see the Materials and methods section) was used to obtain cell monolayers composed of densely-packed cells (to favour intercellular contacts) in a reduced fraction of available space. Under this condition the cultures could be exposed to high volumes of growth medium and thus maintained for long time periods without perturbations. **a:** we show a selection of microphotographs where cells were stained with giemsa. White arrows indicate some multi-nucleated giant cells; black arrows show cytoplasmic bridges between cells; the white dashed arrow indicates mononuclear cells possibly deriving from the multiple cytodieresis of giant multi-nucleated cells. **b:** time-dependent increase of ring diameters of sham-irradiated spheroids (light green squares) or of spheroids treated with a dose of 8 Gy (darker green circles).

Confocal microscopy analyses confirmed the morphological alterations observed following treatment with ionizing radiations, such as the presence of giant multinucleated cells and intercellular cytoplasmic bridges (Fig.3). Actin filaments redistributed in irradiated cells: stress fibres were no longer present and actin formed a layer of filaments parallel to the plasma membrane. We also observed the formation of perinuclear actin rings that are likely to constitute the molecular scaffold for the enucleation of mononuclear cells from giant multinucleated cells (Fig.3).

**Fig.3.**
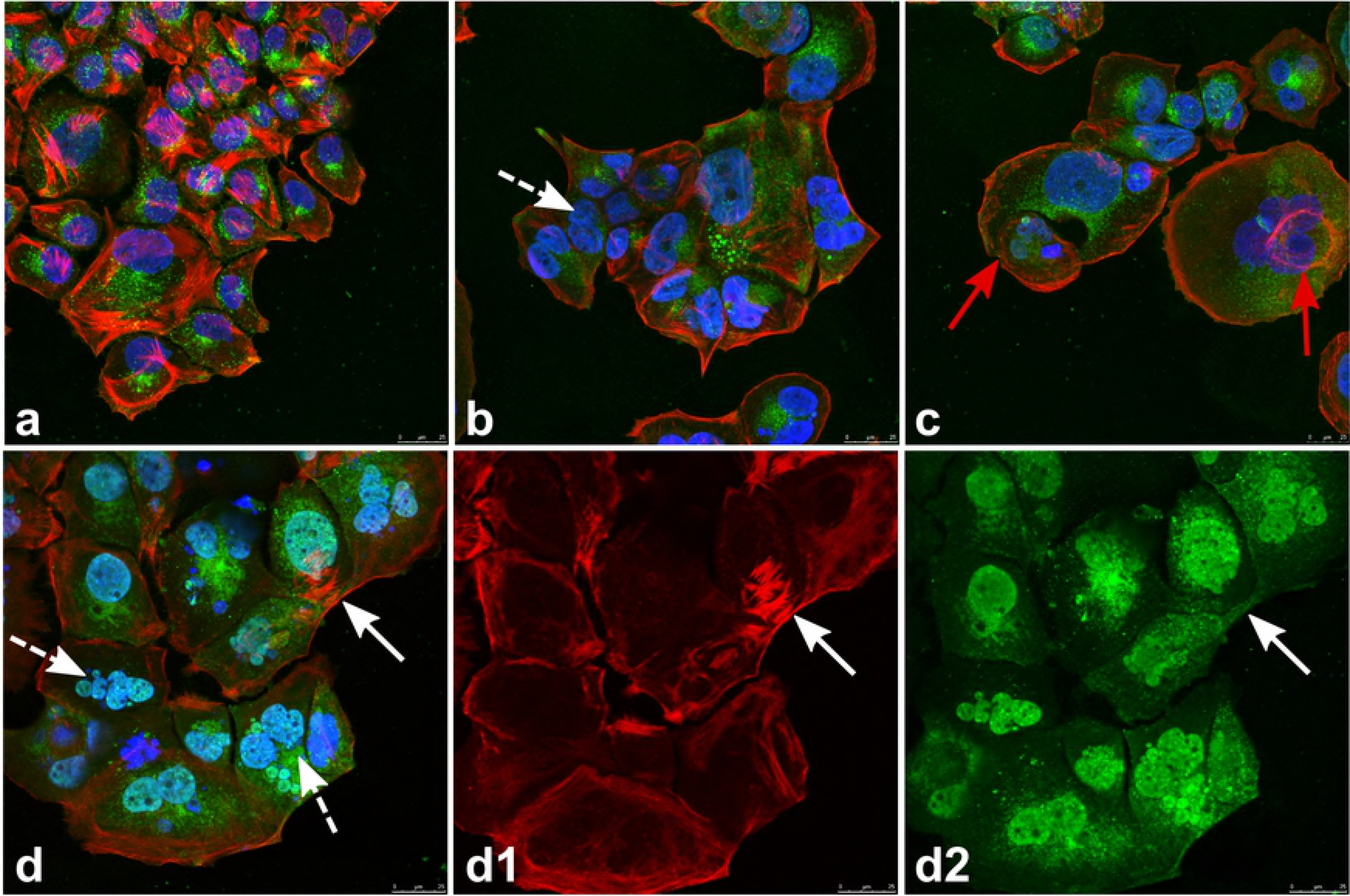
Confocal microscopy of cells in the rings around irradiated (8 Gy) and sham-irradiated tumour spheroids. The cells have been cultured and treated as described in the legend to fig.2 and grown for 480 hours post-treatments. Cells were then fixed, permeabilised and stained with WGA-FITC (green fluorescence; Wheat germ Agglutinin is a protein that binds N-acetyl-D-glucosamine and sialic acid residues ubiquitously present on glycoproteins and glycolipids), rhodamine-phalloidine (red fluorescence; phalloidin binds selectively F-actin filaments), and DAPI (blue fluorescence; the molecule labels the nuclei). **a:** sham-irradiated cells. Stress fibers formed by actin filaments are clearly visible. **b, c.** and **d:** cells irradiated with a dose of 8 Gy. White dashed arrows indicate giant cells with multiple nuclei. In irradiated cells actin filaments redistributed to the plasma membrane and eventually formed perinuclear rings (red arrows). We hypothesize that actin rings could start the scission of new mononuclear cells from multi-nucleated giant cells. The white arrow in **d** indicates a cytoplasmic bridge. **d1 and d2:** images showing only the red and the green fluorescence channels of the microphotograph in **d**. The cytoplasmic bridge (white arrows) indeed connects the cytoplasms of two multi-nucleated cells. The scale bar in all microphotographs is 25 μm long.

Careful inspection of isolated cells in monolayer cultures confirmed complex recovery dynamics comprising events of fusion and division of mononucleated and giant multinucleated cells (see S1 Fig.).

### Role of Syncytin-1 homologous proteins in cell fusion and radioresistance

T47D cells express human endogenous retroviral sequences [17-20] and can produce non infectious viral particles phylogenetically related to mouse mammary tumour viruses [17-20]. Endogenous retroviruses are a family of ancient viruses whose sequences have been, at least in part, conserved during evolution [17, 18]. Indeed, it has been estimated that retroviral DNA sequences occupy up to 8% of the human genome [21, 22]. Retroviruses can express envelope proteins with fusogenic potential, and among the endogenous retroviral proteins Syncytin-1 is the best known molecule for its key role in the formation of the placental tissue syncytiotrophoblast [21, 22].

Since we observed cell fusion between irradiated cells we investigated whether T47D cells could express Syncytin-1 homologous sequences. RT-PCR analysis allowed the amplification of 980 bp-long cDNA fragment showing a 90% of sequence identity with the terminal mRNA portion of Syncitin-1 precursor of *Homo sapiens* (NM_001130925.1, S2 Fig.). Translation of the open reading frame of our cDNA clone provided an amino acid sequence displaying a very high level of identity with the C-terminal regions of Syncitin-1 precursor (NP_001124397.1) (Fig.4). The stretch of 21 amino acids, representing the fusion peptide, and that of 20 amino acids, essential for the fusogenic function, were highly conserved (Fig.4). Thus, T47D mammary carcinoma cells expressed a Syncitin-1 homologous protein (SyHP).

**Fig.4.**
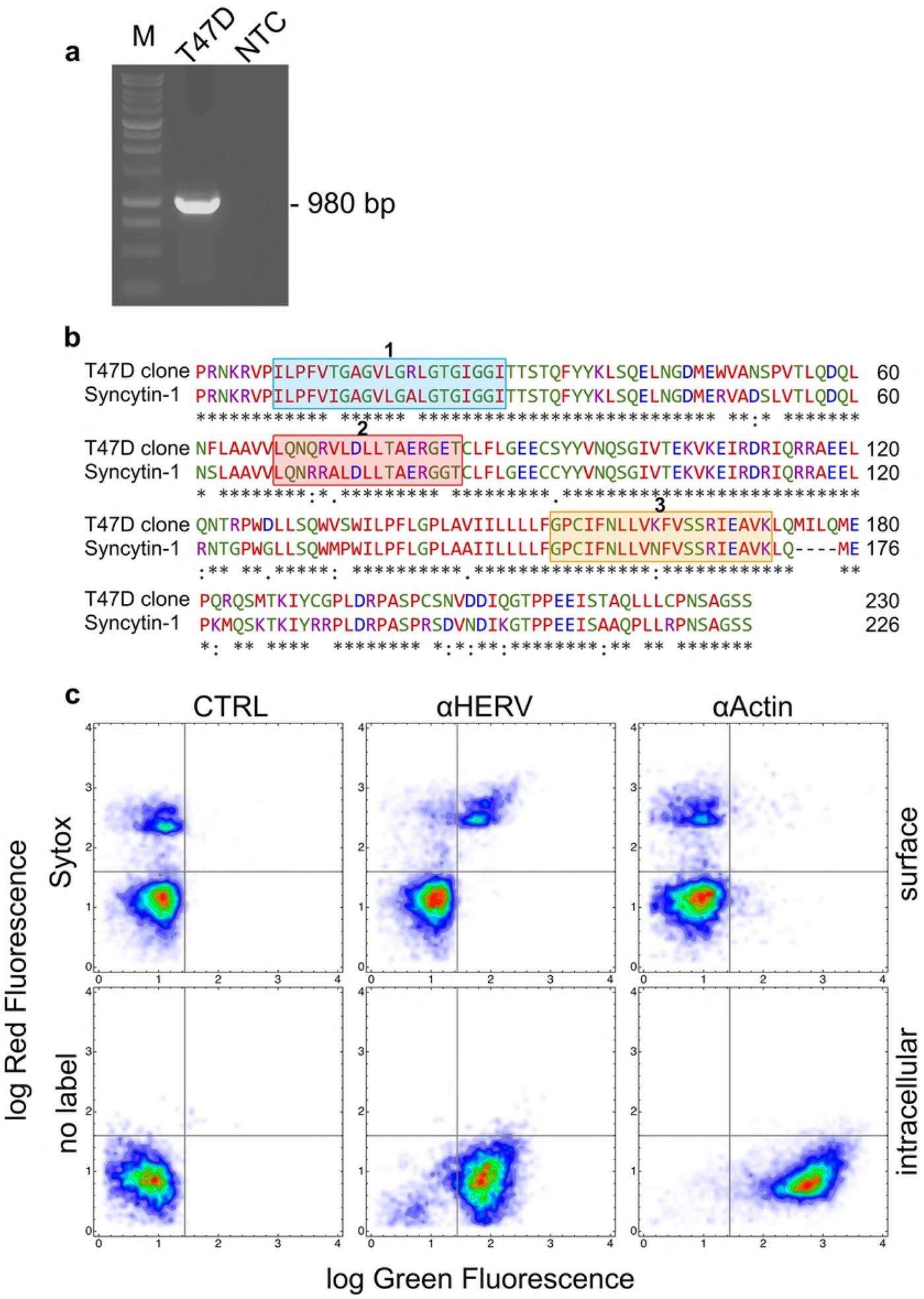
Expression of Syncytin-1 homologous DNA and protein sequences in T47D cells. a: gel elecrophoresis of the DNA fragment amplified by RT-PCR in extracts of T47D cells. The amplicon of 980 bp was obtained with a primer pair designed on the terminal 3′ coding region and a portion of the 3′ untranslated egion of the Syncytin-1 (AF208161) transcript. M: molecular size marker; T47D: fragment amplified in T47D cells; NTC: no template control. The full-length gel is presented in S3 Fig. **b:** the deduced aminoacid sequence of the translatable portion of the amplifed fragment in **a** was aligned against the human Syncytin-1 protein sequence using the Clustal Omega software in search of homologies. The blue box 1 shows the known fusogenic peptide of Syncytin-1 [22] whereas the orange box 3 the domain of Syncytin-1 that is essential for its fusogenic activity [22]. The red box 2 shows the immunosuppressive peptide of Syncytin-1 [22]. T47 cells express a protein (SyHP) with elevated homology for Syncytin-1 **c:** two dimensional flow cytometry investigation of SyHP expression in T47D cells. We used an anti-HERV antibody (aHERV) raised against a central peptide of Syncytin-1. Anti-Actin (aActin) antibodies were used as control. Bound antibodies were revealed with secondary antibodies conjugated to FITC (green fluorescence). Non permeabilised cells were also labelled with the vital fluorescent dye Sytox (red fluorescence) that labels dying and dead cells. Nearly all cells showed intracellular expression of SyHP. SyHP translocated to the plasma membrane of Sytox+ cells.

We also investigated the expression of SyHP by flow cytometry using antibodies raised against a peptide corresponding to the central region of Syncytin-1 (anti-HERV antibodies). Virtually all cells showed cytoplasmic expression of SyHP (Fig.4). Interestingly, we observed surface expression of SyHP in cells labeled with the vital fluorescent dye Sytox, i.e. in cells committed to death or actually dead.

To verify whether dead cells could vehicle fusogenic signals to living cells through the expression of SyHP, we added dead cells isolated from irradiated cultures to non-irradiated cell cultures. Since in crowded cell cultures it was difficult to identify and quantify syncytia, we measured the number of cytoplasmic bridges that clearly correlated with cell fusion (see also the confocal microscopy images shown in Fig.3). Irradiated dead cell bodies induced the formation of cytoplasmic bridges between and among living cells in a dose-dependent manner. Addition of anti-HERV antibodies significantly reduced the number of cytoplasmic bridges (Fig.5). Thus, dead cell bodies delivered fusogenic signals to living cells through the surface expression of SyHP that could be inhibited by anti-HERV antibodies.

**Fig.5.**
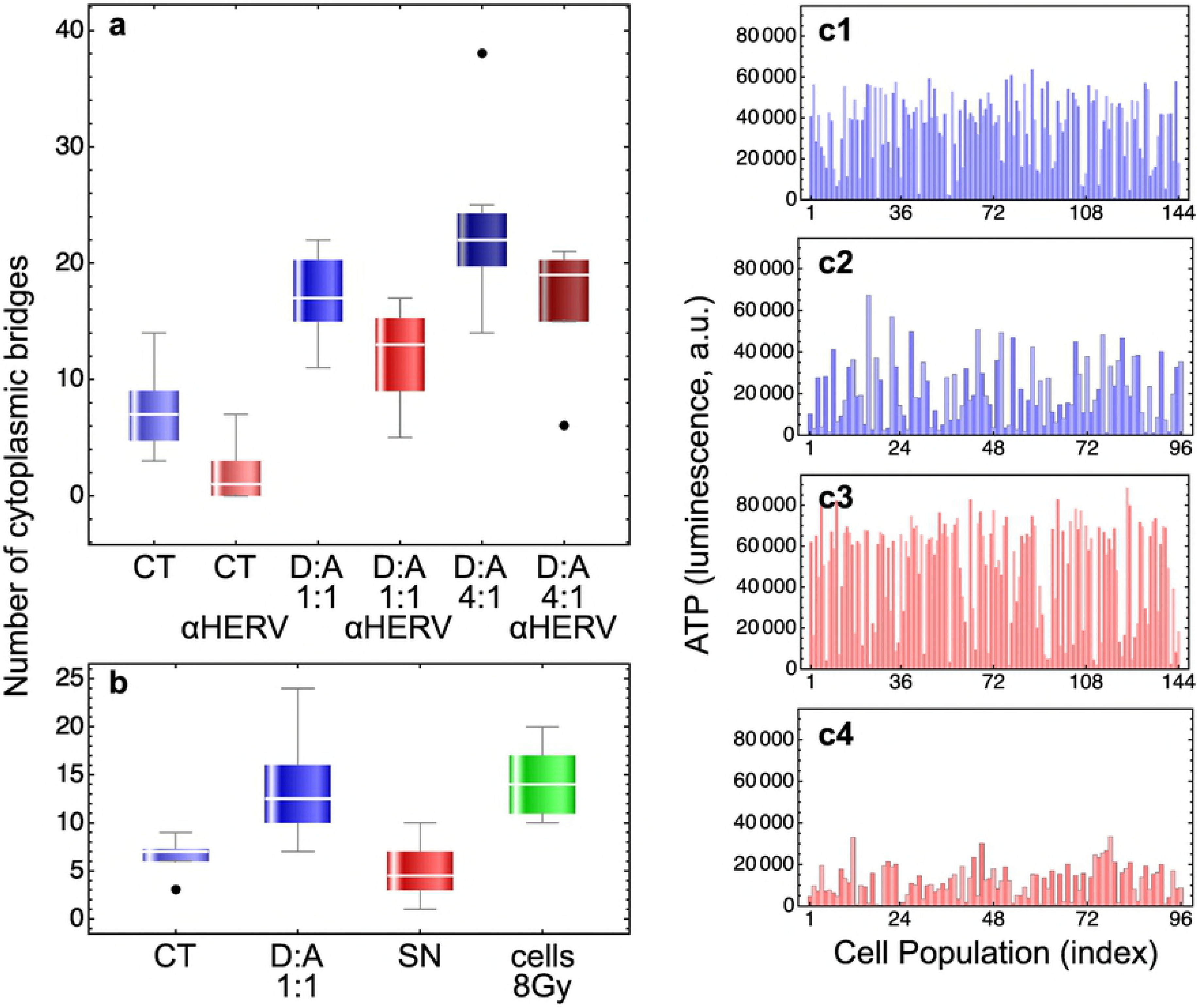
Biological activity of anti-HERV antibodies. a: 10^5^ cells were seeded into the wells of 6-wells culture plates in a final volume of 2 ml of growth medium. Where indicated, we added dead cell bodies (D) that were isolated after 48 hours from the culture supernatants of irradiated (8 Gy) confluent cultures. Cells were scored as dead using the vital dye trypan blue (100%), and were added either at 1:1 or 4:1 ratio with non irradiated alive cells (A). Eventually, we added anti-HERV (αHERV) antibodies diluted 1:100 in growth medium. The batch solutions of anti-HERV antibodies contain 0.09% NaN3 (in PBS) as preservative. Since sodium azide is toxic for cells, we treated cell cultures free from anti-HERV antibodies with 1:100 of 0.09% NaN3 in PBS for control. After 144 hours we counted the cytoplasmic bridges at the microscope from at least 10 randomly selected microphotographs. Data were subjected to two-ways ANOVA analysis, first factor being the addition of dead cells and the second factor the treatment with anti-HERV antibodies. Treatment with dead cells significantly increased the number of observed cytoplasmic bridges *(df=2,* F=81.86, p~10^-17^) whereas anti-HERV antibodies significantly reduced their number *(df=1,* F=55.8, p~10^-10^). No interactions between the two factors were observed *(df=2, F=0.08,* p=0.92). **b:** experiments were carried out as described above. Here, however, we treated the cells also with cell-free supernatants (SN) from irradiated cultures. SN were obtained by centrifugation and filtration (0.22 μm porous filters) of supernatants containing dead cell bodies from irradiated cultures. The number of cell bridges was determined also for cells in standard cultures irradiated with a dose of 8 Gy. Data were subjected to one-way ANOVA analysis followed by Bonferroni’s and Tukey’s post-hoc tests. Overall the differences, where observed (CT or SN vs. D:A, CT or SN vs. cells 8 Gy), were statistically significant *(df=3,* F=17.3, p~10^-7^). Importantly, treatment with cell-free supernatants did not increase the number of cytoplasmic bridges, demonstrating that the effects were caused by dead cells. **c:** we show a few examples of the experiments reported in Table 1. Cell populations were grown in F-bottom wells (**c1** and **c2**) or V-bottom wells (**c3** and **c4**) by seeding an initial number of 3500 (F-bottom) or 2500 (V-bottom) cells in 100 μl of growth medium. After 24 hours cell population were treated with a dose of 8 Gy of ionizing radiations in the presence (**c2** and **c4**) or in the absence (**c1** and **c3**) of anti-HERV antibodies at 1:500 final dilution. NaN3, 1:500 from a 0.09% stock solution in PBS, was also added for control (**c1** and **c3**). After ~35 days ATP levels in each population was determined by the luciferine-luciferase assay. Threshold ATP levels (~9500 luminescence units) were determined with cells treated with a lethal radiation dose of 20 Gy. Cell populations with an ATP level above the threshold were scored as surviving treatment (see also Table 1).

To test whether cell fusion could have a role in promoting cell radioresistance, we exposed several cell populations to ionizing radiations in the absence or presence of anti-HERV antibodies. Treatments with anti-HERV antibodies reversed the radioresistance phenotype and cells became sensitive again to radiation (Fig.4 and Table 1); antibodies also abrogated the radioresistance observed when cell populations were grown in conical-bottom wells (Table 1).

**Table 1.**
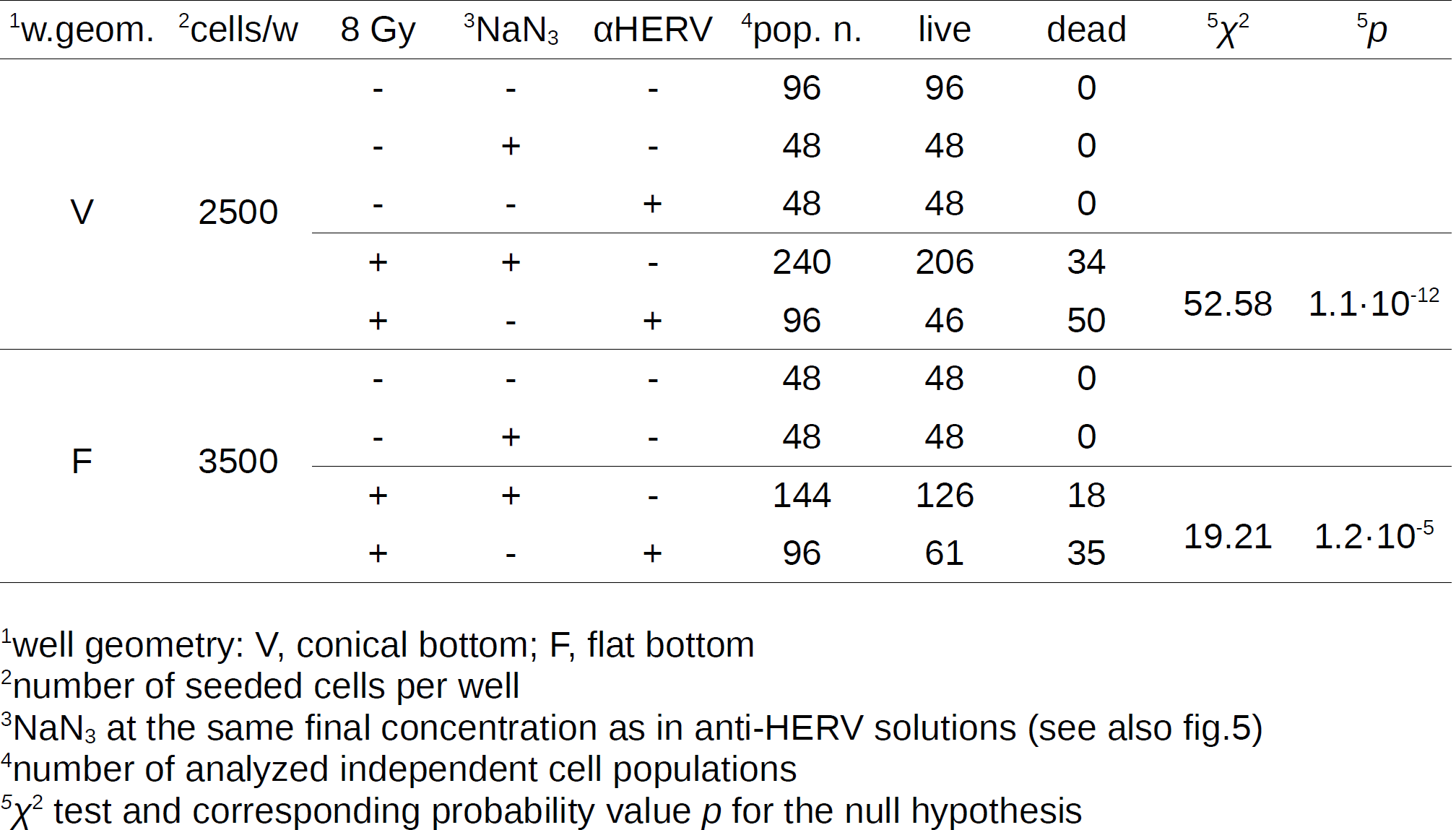
Sensitivity of T47D cells to radiation and aHERV treatments

## Discussion

Polyploidy associated with genotoxic stress, particularly radiation, is well documented [23-26]. Originally, the induction of polyploidy was viewed as an irreversible process leading to cell death, but now there are indications that this might not be necessarily the final fate of the cells (for a review see [26]). Indeed, for a small fraction of cells, polyploidy can be reversible and can provide an escape mechanism to genotoxic insult [23-26]. It has been reported that polyploidization occurs in 10-50% of the cells and that extended cell death leaves only 10-20% of polyploid giant cells alive [26]. Of these the vast majority slowly senesces leaving a small fraction of cells that can de-polyploidize to re-acquire self renewing potential [26].

Our data are consistent with this complex biological model of cell survival from genotoxic stress. The data, however, allow us to extend this model by including fusion events that take place at the cell population level and that result in significantly higher tumour cell survival. We indeed observed the formation of giant cells with multiple nuclei, but we also observed a quite complex dynamic behaviour of the cells during recovery when both giant multinucleated and individual cells were found to fuse and divide. It was not clear whether the formation of multiple nuclei in giant T47D cells derived from endoduplication of nuclear material or from cell fusion. Since events of cell fusion and division could be observed it is likely that both processes might occur in these cells. Experiments carried out with anti-HERV antibodies, nonetheless, show that envelop proteins homologous to Syncytin-1 from endogenous retroviruses are primarily involved in cell fusion and resistance to radiation treatments of T47D breast carcinoma cells.

Endogenous retroviruses are abundant in mammalian genomes and their sequences have been shown to regulate innate immunity [27]. The envelope glycoprotein of retroviruses is essential for viral entry and mediates the fusion between the viral and the plasma membranes. The protein is produced as a single precursor polypeptide chain which is then cleaved by cellular proteases to form the surface (SU) and the transmembrane (TM) subunits. The two subunits remain associated as heterodimers which are further assembled into protein trimers at the surface of the envelope membrane [22]. SU mediates receptor recognition whereas TM cell fusion through refolding of the subunits and unmasking of its fusogenic peptide [22]. Endogenous envelop proteins may have kept, lost or modified their original function during evolution. The most studied case is that of Syncytin-1: knockout of syncytin genes in mice provide clear evidence that they are absolutely required for placenta development and embryo survival by inducing fusion of syncytial cells at the foetal-maternal interface [22].

T47D cells can express endogenous retroviral sequences and produce viral-like particles [17-20]. These particles are considered not infectious [19], and indeed in our experiments the supernatants of irradiated cultures did not show any fusogenic activity. Interestingly, also the immunosuppressive peptide [22] of Syncytin-1 is conserved in SyHP expressed by T47D cells. It would be interesting to explore whether this peptide could mediate evasion of breast cancers from immune surveillance.

In conclusion, we report on a complex mechanism of tumour cell radioresistance that involves endogenous retroviral proteins and cell fusion. These mechanisms can only be observed at the cell population level and measured using appropriate experimental approaches. Measuring the effects of radiation treatments by focusing on clonogenic cells only may lead to overestimate the sensitivity of tumours to radiotherapy. Our theoretical and experimental approach can overcome the limitations of clonogenic assays.

## Supporting Information

**S1 Fig. Cytological analysis of T47D monolayers after irradiation with 8Gy.** Typical examples of microphotographs taken at the indicated post-irradiation times. The microphotograph at 0 hours was taken within 15 min from irradiation. A group of cells is clearly visible. After 72 hours post-irradiation these cells showed an enlarged cell volume (black arrow). At 72 hours some of the individual cell bodies were visible but at later times (96 and 128 hours) cell fusion and the formation of giant cells with multiple nuclei was evident (black arrow). At 144 hours (black arrow), however, a division event took place that produced two giant multinucleated cells. One of these giant cells shrank at later times (white arrows) but the other apparently fused again with neighbour giant cells (black arrows) to form a lump of cells (264 hours, black arrow). Expulsion of cell material from the cluster was then observed after 288 hours post-irradiation. In all panels the reference bar is 100 pm long.

**S2 Fig. Sequence analysis of the 980 bp-long cDNA fragment amplified by RT-PCR in T47D cell extracts.** The amplified DNA fragment encompasses a region of 683 nt of the 3’ terminal coding region and a portion 276 nt of the 3’untranslated region of the Syncytin-1 (AF208161) transcript. Alignment between the two DNA sequences was carried out using Clustal Omega software hosted at https://www.ebi.ac.uk.

**S3 Fig. Gel elecrophoresis of the DNA fragment amplified by RT-PCR in extracts of T47D cells.** Picture of the full-length gel from which we cropped the image reported in Fig.4A in the main text.

Contributions
RC, MS, and EM conceived the project. RC and MS designed and performed the experiments with tumour cells and spheroids. BM designed and performed molecular biology experiments. AB designed and performed morphological experiments and image analyses. RC and EM developed mathematical models. All authors discussed the results and wrote the manuscript.

## Notes

**Competing interests:** the authors declare no competing interests

